# Molecular basis of Progressive Familial Intrahepatic Cholestasis 3. A proteomics study

**DOI:** 10.1101/2023.06.15.545058

**Authors:** Laura Guerrero, Lorena Carmona-Rodríguez, Fátima Millhano Santos, Sergio Ciordia, Luiz Stark, Loreto Hierro, David Vicent, Fernando J. Corrales

**Author notes:** To whom correspondence should be addressed Corresponding Author, Fernando J. Corrales, Ph.D., Functional Proteomics Laboratory, National Center for Biotechnology, CNB-CSIC, Darwin, 3, 28049 Madrid, Spain. **Authors’ contributions:** Experiments and procedures: LG, LCR, FMS, SC, LS, DV, LH and FJC; patient handling: LH; concept and design: DV and FJC; supervision: FJC; writing of article: FJC and LG. **Ethics approval statement** The animal study protocol was approved by the *Comunidad de Madrid* (PROEX 845/2019) and the CNB Ethics Committees in strict accordance with the Spanish and European Union laws and regulations concerning the care and use of laboratory animals. The human study was approved by the Clinical Research Ethics Committee of Hospital Universitario La Paz (HULP PI-3252) in agreement with the principles stated in the Declaration of Helsinki. **Patient consent statement:** all participants provided informed consent as established by the Hospital Universitario La Paz. **Permission to reproduce material from other sources:** Not applicable. **Availability of data and materials** The dataset used and/or analyzed during the current study is available from the corresponding author on reasonable request. **Supplementary Materials:** supplementary figures (1-3), supplementary tables (1-5), supplementary methods.

## Abstract

**Background and aims:** Progressive familiar intrahepatic cholestasis type 3 (PFIC3) is a severe rare liver disease which affects between 1/50,000 to 1/100,000 children. In physiological conditions, bile is produced by the liver and stored in the gallbladder, then it flows to the small intestine to play its role in fat digestion. To prevent tissue damage, bile acids are kept into phospholipid micelles. Mutations in phosphatidyl choline transporter ABCB4 (*MDR3*) lead to intrahepatic accumulation of free bile acids that results in liver damage. PFIC3 onset occurs usually at early ages, progress rapidly and the prognosis is poor. Currently, besides the palliative use of ursodeoxycholate, the only available treatment for this disease is liver transplantation, which is really challenging for short-aged patients.

**Methods:** To gain insight into the pathogenesis of PFIC3 we have performed an integrated proteomics and phosphoproteomics study in human liver samples to then validate the emerging functional hypotheses in a PFIC3 murine model.

**Results:** We identified 6,246 protein groups, 324 proteins among them showing differential expression between control and PFIC3. The phosphoproteomic analysis allowed the identification of 5,090 phosphopeptides, from which 215 corresponding to 157 protein groups, were differentially phosphorylated in PFIC3, including MDR3. Regulation of essential cellular processes and structures, such as inflammation, metabolic reprograming, cytoskeleton and extracellular matrix remodeling and cell proliferation were identified as main drivers of the disease.

**Conclusion:** Our results provide a strong molecular background that significantly contributes to a better understanding of PFIC3 and provides new concepts that might prove useful in the clinical management of patients.

**Lay Summary/Key Points:** PFIC3 is a rare disease that affect 1/50,000 to 1/100,000 children that present severe symptoms in the first years of life and have scarce therapeutic options. We identified a panel of proteins that recapitulate some of the main cellular processes associated to the progression of liver injury in PFIC3 patients and suggest alternative treatment options.

## Introduction

Cholestasis is a pathological condition of the liver that results from impaired formation or flow of bile. This defect can occur anywhere between the basolateral membrane of the hepatocytes and the ampulla de Vater, leading to intrahepatic or extrahepatic cholestasis. A consequence of the altered bile dynamics is the accumulation of biliary components that induce a cellular damage to the liver parenchyma, the biliary tree and the circulation of harmful compounds, including bile acids. The clinics of cholestasis is diverse, from mild to severe symptoms, such as prutritus, fatigue, abdominal pain, jaundice, elevation of aminotransferases and in some cases GGT and alkaline phosphatase. Its presentation, either acute or chronic, leads to impaired hepatobiliary function, progression of different cholestatic syndromes, fibrosis, cirrhosis and hepatocellular carcinoma (HCC) or cholangiocarcinoma, in the most severe cases. In adults, cholestasis spans a wide array of etiologies: genetic, drug-induced liver injuries, pregnancy, sepsis, biliary obstruction and autoimmune. In children, relevant pathophysiology arises from genetic cholestasis syndromes (Progressive Familial Intrahepatic Cholestasis, Alagille syndrome) and biliary atresia ^1^.

Progressive familial intrahepatic cholestasis (PFIC) represents a group of autosomal recessive disorders that affects 1 in 50,000 to 100,000 newborns ^2^. The disease is presented during the childhood as an intrahepatic cholestasis derived from defects of bile acid metabolism and transport^3^. The phenotype of PFIC range from benign recurrent cholestasis, to a severe syndrome that might lead to liver failure and cancer within the first ten years of life. Three different types of PFIC have been identified according to the responsible mutated genes^4-6^. PFIC1 associates to the mutation of ATP8B1, which encodes the amino-phospholipid flippase FIC1; PFIC2 involves mutations on ABCB11 gene that encodes for the bile salt export pump BSEP; and PFIC3 that results from mutations on ABCB4 gene encoding the phosphatidyl choline floppase multidrug resistance P-glycoprotein (MDR3).

ABCB4 gene is located on chromosome 7q21, consists of 27 coding exons spanning 74 kb^7^ and expresses mainly in hepatocytes^8^ where the encoded protein MDR3 is trafficked to the apical membranes to form the boundary to bile canaliculi. The protein has two transmembrane α-helical domains that adopt a collapsed conformation at the level of the lipid bilayer, which contain the substrate binding pocket, and two ATP binding cytosolic domains^9^. Although there are different hypotheses about the precise mechanism by which MDR3 drives the translocation of phosphatidyl choline (PC) across the membrane, it appears to involve conformational changes driven by the ATPase activity^10,11^. PC is an essential component of bile involved in the formation of micelles, which keep the cholesterol and insoluble bile salts in soluble form, protecting the cholangiocytes from damage. Currently, about 300 ABCB4 variants associated with a progressive cholestatic liver disease have been described. Attending to the functional effects induced into the encoded MDR3, ABCB4 variants have been classified into 5 classes: I, nonsense mutations preventing MDR3 synthesis; II, missense variants that impede protein maturation; III, missense mutants with compromised activity; IV, variants leading to unstable proteins; V, unknown pathogenicity^12^. A defective transport of PC across the bile canaliculi results in a free bile salts-mediated cellular injury as well as in the formation of stones of crystallized cholesterol that obstruct small bile ducts, damaging liver structures. Although cholestatic liver disease may develop in pediatric subjects resulting from partial defects on MDR3 activity^13^, heterozygosity commonly associates with mild manifestations, while homozygous individuals present severe and potentially life-threatening complications^14^. Different to other PFIC, PFIC3 can be presented in adulthood but in the early onset, pruritus, hepatosplenomegaly, portal hypertension and variceal bleeding, acholic stools, stunted growth, jaundice, reduced bone density and learning disabilities have been associated with MDR3 impairment^15,16^. There are different therapeutic options currently available that are selected according to the severity of the disease and the nature of the molecular defect. Ursodeoxycholate treatment have shown effective in up to 70% of cases maintaining some residual MDR3 activity and PC secretion, chemical chaperones could improve MDR3 trafficking when the shuttling to the plasma membrane is compromised, pruritus can also be improved with rifampicin. However, while expecting the consolidation of emerging substitutive treatments based by gene therapy or RNA technology approaches, liver transplant is the only known curative treatment for severe PFIC3^14^. However, the availability of organs for pediatric cases is limited and therefore any intervention that may contribute to stabilize patients by reducing the progression pace of the disease would improve their clinical expectancies. In this regard, the systematic analysis of the molecular basis of PFIC3 progression might provide useful information to develop new strategies for a better management of PFIC3 patients. In this study, we have performed a combined analysis of the proteome and phosphoproteome of liver samples from control individuals and PFIC3 patients. Our results provide a detailed understanding of the main cellular processes regulated in cholestatic hepatocytes, lead to the identification of some of the drivers of PFIC3 progression and, based on this information, suggest strategies that may pave the way for the development of improved clinical interventions.

## Materials and methods

### Biological samples

Human tissue samples. Liver tissue fragments from the explanted livers of PFIC3 patients (N=4) and from the liver graft of living donors (N=4) were obtained at the time of liver transplantation and stored frozen in RNAlater solution (Invitrogen). ABCB4 pathogenic mutations were documented in all PFIC3 by genetic analysis. Liver transplant were performed between 6 and 9 years old. All surgical procedures were conducted in Hospital Universitario La Paz. Liver samples were kept at the institutional biobank. Written informed consent was obtained from patient’s legal guardians and graft donors. Study protocols conformed to the principles stated in the Declaration of Helsinki and as such they were approved by the Clinical Research Ethics Committee of Hospital Universitario La Paz.

*Mdr2* -/- and wild type mice liver. Age: 5 months (WT: n=3; *Mdr2*: -/- n=3), 9 months (WT: n=3; *Mdr2* -/-: n=3), and 17 months (WT: n=3; *Mdr2* -/- non-tumor: n=3; *Mdr2* -/- tumor: n=3),

### Lysis and protein extraction

Liver samples (human and mice) were disrupted mechanically using a Potter–Elvehjem homogenizer in lysis buffer containing 5% SDS (Sodium dodecyl sulfate) (Sigma-Aldrich), 100 mM triethylamonium bicarbonate (Thermo Fisher Scientific) and a protease/phosphatase inhibitor cocktail (Thermo Fisher Scientific). For removing DNA, samples were sonicated by micro tip probe ultrasonication for 1 min on UP50H ultrasonic lab homogenizer (Hielscher Ultrasonics). After centrifugation at 10,000 × *g* for 5 min, the protein concentration of the saved supernatant was measured using the Pierce 660-nm Protein Assay adding IDCR (Ion detergent compatibility reagent) (ThermoFisher Scientific). Supernatant containing the proteins were stored at -80ºC until further analysis by LC-MS and western blot.

### Liquid chromatography and mass spectrometry analysis (LC-ESI-MS/MS)

Liver proteins were reduced and alkylated by adding 5 mM TCEP (tris(2-carboxyethyl) phosphine) and 10 mM chloroacetamide for 30 minutes at 60ºC before digestion into peptides. Protein digestion was performed using S-Trap filter (Protifi, Huntington, NY, USA), according to the protocol. We digested 80μg of protein and added trypsin (Thermo-Fisher Scientific) in a ratio 1:20 (trypsin:protein).

The resulting peptides were subsequently labelled using TMT-11plex Isobaric Mass Tagging Kit (Thermo Scientific, Rockford, IL, USA) according to the manufacturer’s instructions as indicated in supplementary materials and methods. Labelled peptides were used for the analysis of the global proteome and the phosphoproteome.

For deep analysis of the global proteome, high pH reversed-phase peptide prefractionation was performed with sulfonated divinylbenzene (CDS Empore™ SDB-RPS, Sigma-Aldrich) using a step gradient of increasing acetonitrile concentrations (0-60% ACN) in a high pH elution solution (10 mM ammonium formate). Eluted peptides were collected into 10 different fractions by centrifugation at 850 xg during 3 minutes. The peptide fractions were dried and stored at −20 °C until the LC−MS analysis.

In parallel, phosphopeptide enrichment was performed using TiO_2_ columns and cleaning of the sample was performed with Oligo R3 reverse-phase before LC-MS analysis^1^.

Peptides were solubilized in 2% acetonitrile (ACN) and 0.1% formic acid (FA), then peptide concentration was determined by Qubit 2.0 Fluorometric Quantitation (Thermo Fisher Scientific). 1 μg aliquot of each fraction in an injection volume of 5 μl was analyzed by nano LC ESI-MS/MS (Liquid Chromatography Electrospray Ionization Tandem Mass Spectrometry) in an Ultimate 3000 nano HPLC system (Thermo Fisher Scientific) coupled to an Orbitrap Exploris 240 (Thermo Fisher Scientific). Peptides were loaded onto a 50 cm × 75 μm Easy-spray PepMap C18 analytical column at 45°C and were separated at a flow rate of 300 nL/min using a 120 min gradient ranging from 2 % to 95 % mobile phase B (mobile phase A: 0.1% FA; mobile phase B: 80 % ACN in 0.1% FA).

Data acquisition was performed using a data-dependent top-20 method, in full scan positive mode, as indicated in supplementary methods.

### Proteomics data analysis

Details are provided into the supplementary material. In brief, raw files were processed using Proteome Discoverer (PD) version 2.4 (Thermo Fisher Scientific). MS2 spectra were searched using four search engines and a target/decoy database from the human proteome at Uniprot Knowledgebase (9606rev_20210219). The peptide precursor mass tolerance was 10 ppm, and MS/MS tolerance was 0.02 Da. The false discovery rate (FDR) for proteins, peptides, and peptide spectral matches (PSMs) peptides were kept at 1%.

The quantification values for proteins were calculated using the abundance of total peptide.

To calculate the p-values and adjusted p-values we used a t-test. Significance was considered as adjusted p-value >= 0.05. A list of differentially expressed and phosphorylated proteins was used to perform an Ingenuity Pathway Analysis (IPA).

### Monitoring One-carbon metabolism by targeted proteomics (MRM)

Details are provided in the supplementary methods. For monitoring One-carbon metabolism in liver samples, 12 participating enzymes were monitored by multiple reaction monitoring (MRM) targeted proteomics ^17^. For MRM analyses, 1 μg of peptide analyzed on a 5500 QTRAP triple-quadrupole mass spectrometer. Raw MRM data files were analyzed with Skyline software (v21.2.0.568). Statistical analysis and graphical representation of data were performed using GraphPad Prism Software v9.3.1 was used. Statistical significance was considered for p-value <0.05 according to t-test analysis results.

### Western blot

20 μg of protein in loading buffer (10% glycerol, 2 % SDS, 5% β mercaptoethanol, 0.062 M Tris-HCl, 0.002 % bromophenol blue) were analyzed by electrophoresis in 12% acrylamide at constant amperage (20 mA). The proteins were transferred to 0.2 μm nitrocellulose membranes, which were blocked with 1 × TBS/0.05% Tween containing 5% milk or 5% bovine seroalbumin (BSA). Membranes were then incubated with primary antibodies at 4 °C overnight in 1 × TBS/0.05% Tween containing 5% BSA/milk, and then in horseradish peroxidase (HRP)-conjugated secondary antibodies (Dako) at room temperature for 1 hour and then developed with WesternBright ECL HRP substrate (Advansta). The list of used antibodies is indicated in supplementary materials and methods. Statistical analysis of the results was performed using R v.4.1.3. For comparisons between two groups, the t-test function of R Base package was used. Differences were considered significant when p value <0.05.

### RT-qPCR

For RNA extraction, liver tissue samples were homogenized using a Pellet Pestle in buffer provided by RNeasy kit (Qiagen) according to manufacturer instructions. RNA concentration was determined using a Nanodrop ND-1000 (Thermo Fisher Scientific). 1 μg of RNA of each sample was retrotranscribed using the Applied Biosystems Kit RNA-to-cDNA. Quantitative PCR was then performed using EvaGreen Master Mix (Solis BioDyne) detected by the QuantStudio 5 and analyzed using QuantStudio Software v1.5.2 (Applied Biosystems). Primer sequences used for qRT-PCR are shown in supplementary materials and methods. Results were normalized to the expression of GAPDH and presented as fold induction with respect to the wild type samples according to the 2^-ΔΔCT^ method. Statistical analysis of the results was performed using R v.4.1.3. For comparisons between two groups, the t-test function of R Base package was used. Differences were considered significant when p value <0.05.

## Results

### Identification of regulated proteins in the liver of PFIC3 patients

To shed light into the molecular principles driving the progression of PFIC3, we have combined a differential proteomic and phosphoproteomic analysis comparing liver samples from control individuals and PFIC3 patients. Subjects with four independent ABCB4 mutations that encode four different protein variants were analyzed: p.G68R (homozygote), p.R1187X + c.3633+2 (intronic, involves a splicing site) (compound heterozygote), p.D459H+P479L (compound heterozygote) and p.S436N (homozygote). While G68 locates at one of the transmembrane helices of MDR3, with an undetermined effect on the protein structure/function, the rest of the amino-acid substitutions occur in one of the two ATP binding domains, likely compromising the binding or hydrolysis of the nucleotide. One of the open questions to be addressed is whether the different ABCB4 mutations analyzed here may have different effect on MDR3 activity and therefore may induce alternative mechanisms of MDR3-associated cholestasis.

Overall, 6,625 proteins corresponding to 5,808 protein groups were identified with an FDR≤1% and quantified (q≤1%) resulting in 292 differential proteins, from which 166 and 126 were up- and down-regulated respectively. Upon TiO2 enrichment, 5,731 peptide groups were identified, corresponding to 1851 protein groups (FDR≤1%), revealing 4,345 phosphorylation sites with more than 90% confidence. Among them, 215 displayed a differential profile, 80 up-regulated and 135 down-regulated PFIC3 livers (Figure 1 A and B). The overlap between the panels of regulated proteins by change in the abundance or phosphorylation is negligible, indicating that the differential phosphorylation events cannot be explained by protein abundance changes.

**Figure 1.**
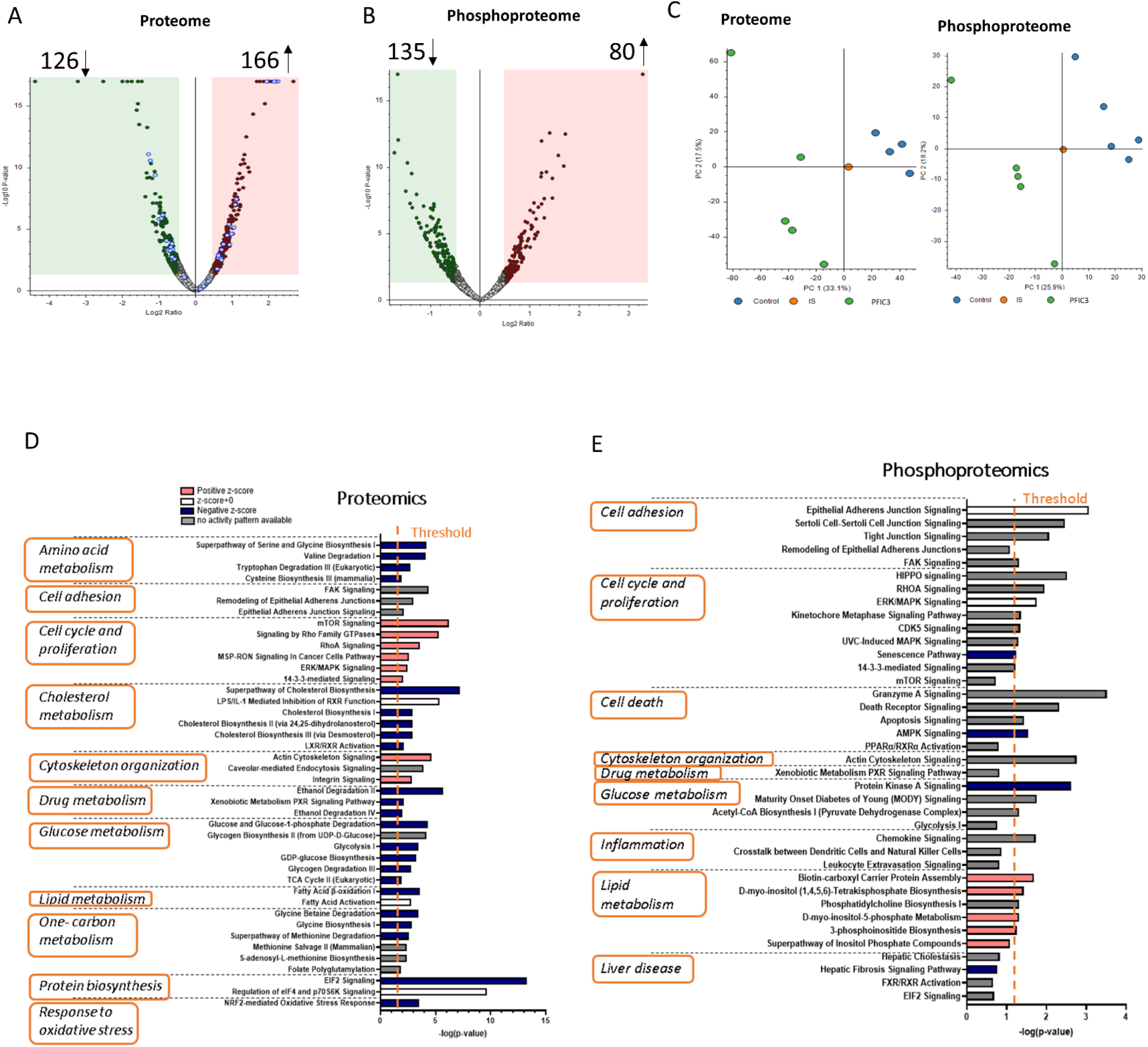
Comparative shotgun proteomics and phospho-proteomics analysis of normal vs PFIC3 liver samples. A and B, volcano plot (−log10 [*p*-value] and log2 [fold-change]) of the proteins and phosphoproteins identified in normal and PFIC3 liver samples. Proteins and phospho-proteins are represented as black dots (p-value < 0.05), red dots (p-value < 0.05; log_2_ fold change < -1) and green dots (p-value< 0.05; log_2_ fold change > 1). C, PCA analysis of both proteomics and phospho-proteomics data set showing the segregation of normal vs PFIC samples. IS correspond to the internal standard that contains equal amounts of each individual sample. D, pathways that are predicted to be regulated from the panel of differential proteins in PFIC3 vs normal samples. E, pathways that are predicted to be regulated from the panel of differentially phosphorylated proteins in PFIC3 vs normal samples. Ingenuity pathway analysis (IPA) of the differentially represented proteins between normal and PFIC3 livers allowed to identify the top enriched canonical pathways. X-axis indicates the significance (-log p-value) of the functional association that is dependent on the number of genes in a class as well as on the biological relevance. The dashed line marks the statistically significant threshold.

Both differential proteins and phosphoproteins efficiently segregate control and PFIC3 cases, likely providing molecular insights on the cellular process involved in the onset of the cholestasis associated to the MDR3 deficiency (Figure 1C). It is noteworthy that we included two replicates of one of the samples, the compound heterozygote, with the purpose of assessing the reproducibility of our analysis and to discard major biases resulting from the experimental workflow. The technological replicates showed a reasonable overlapping in the PCA analysis, suggesting that the dispersion among the protein profile of the analyzed samples has a biological origin.

To understand the cellular pathways altered in PFIC3, a functional enrichment analysis was conducted using the Ingenuity Pathway Analysis (IPA) software. Despite the limited overlapping between the differential protein and phosphoprotein panels, a common functional landscape was deduced from both protein sets, as could be expected since they represent two regulatory levels of the same pathogenic process (Figure 1 D and E). Among them, metabolic rewiring, inflammation, liver differentiation, proliferation and cell adhesion and cytoskeleton were further investigated using the MDR2-/-mice, a well-accepted model of PFIC3 that involves inflammation, cirrhosis and ultimately HCC, at least in the first stages of disease progression^12^.

Among the quantified proteins, MDR3 showed similar levels in all PFIC3 patients than in control individuals, suggesting the normal synthesis and stability of the protein. However, we cannot rule out the possibility of a defective shuttling of any of the MDR3 variants to the plasma membrane of the hepatocyte. Besides the effect of the amino-acid variants, we describe here a putative regulatory mechanism of MDR3 activity based on phosphorylation (Figure 2A). All variants analyzed showed a marked decrease of S666 and T667 phosphorylation (Figure 2B), which are located in a flexible loop bridging the two ATP binding domains. The assignment of the -LFRHpSpTQKNLK-peptide spectrum was verified with the analogous synthetic peptide containing the two phosphoresidues (Figure 2C). The spatial configuration of this region cannot be deduced from previous structural analysis^9,10^ due to its mobility but for representation purposes we have simulated a potential arrangement using alpha-fold^18^ (Figure 2D). A potential mechanism to regulate MDR3 activity has been already suggested based on the phosphorylation of T34 and additional neighboring residues in a PKC-dependent mechanism^19^.

**Figure 2.**
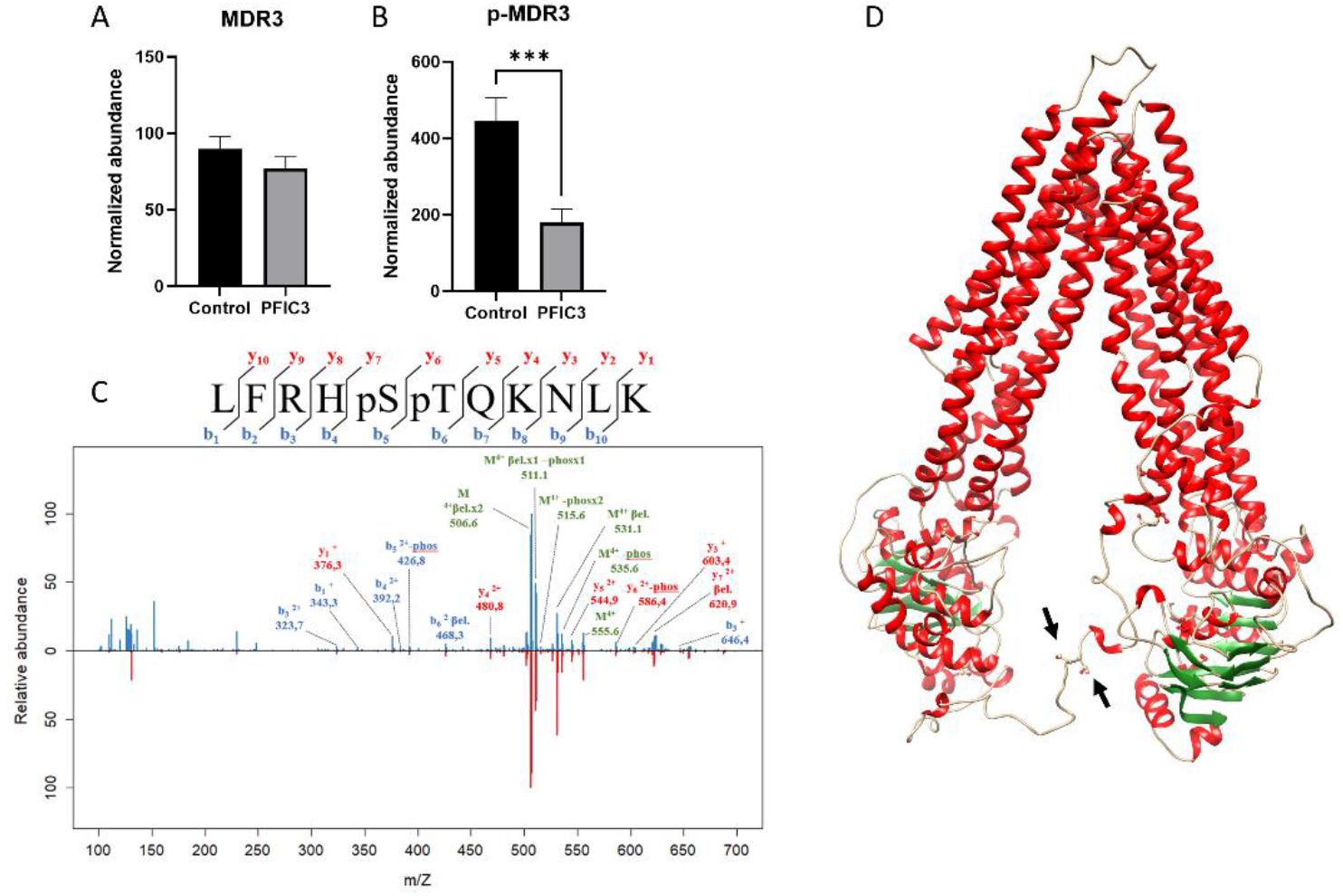
Differential phosphorylation of MDR3 protein in PFIC3 compared to normal livers. A, hepatic levels of MDR3 protein as estimated by mass spectrometry. B, quantitation of the phosphorylation levels of the MDR3 peptide LFRHpSpTQKNLK. C, annotated spectrum of the phosphorylated peptide; the experimental data are in blue and the spectrum resulting from the MS/MS fragmentation of a synthetic peptide containing pS and pT residues at the same positions as the endogenous form is in red. D, structural model of MDR3 based on alpha-fold calculations. The overall structure is based on experimental data (JA Olsen et al Nat Struct Biol 2020) and the theoretical position of the loop connecting the two nucleotide binding domains was predicted.The arrows indicate the position of the pS666 and pT667 residues.

### Inflammation, differentiation and proliferation in PFIC3 livers

Cholangiocytes in the small bile ducts express receptors that allow proliferation and inflammation in response to liver injury^20^. The accumulation of free bile salts due to defective phospholipids in bile to neutralize their detergent effects, results in an injured biliary epithelium and canalicular membrane in PFIC3 patients, leading to intrahepatic cholestasis. Chronic liver diseases including cholestasis, initiate immunoinflammatory responses mediated by a variety of effectors^21,22^, HLA and costimulatory molecules among them^23^. We found that overexpression of HLA I (HLA-B) and HLAII molecules (HLA-DMB, HLA-DPA1, HLA-DPB1 and HLA-DRB3) in PFIC3 livers (Figure 3), underlying the inflammatory response of PFIC3 livers. In agreement, it has been reported that cholangiocytes can express class I HLA under normal conditions but they express HLAII mainly under chronic liver injury^1^. The accumulation of immunoglobulins (IGHG1 and IGHG4) also suggests the activation of the humoral response (Figure 3). To further confirm the chronic inflammation induced by the deficiency of MDR3 in the liver, we measured the levels of TNFa in the liver of MDR2-/-mice. In agreement to previous studies^24^, we observed a progressive accumulation of TNFa in deficient livers (Figure 3B).

**Figure 3.**
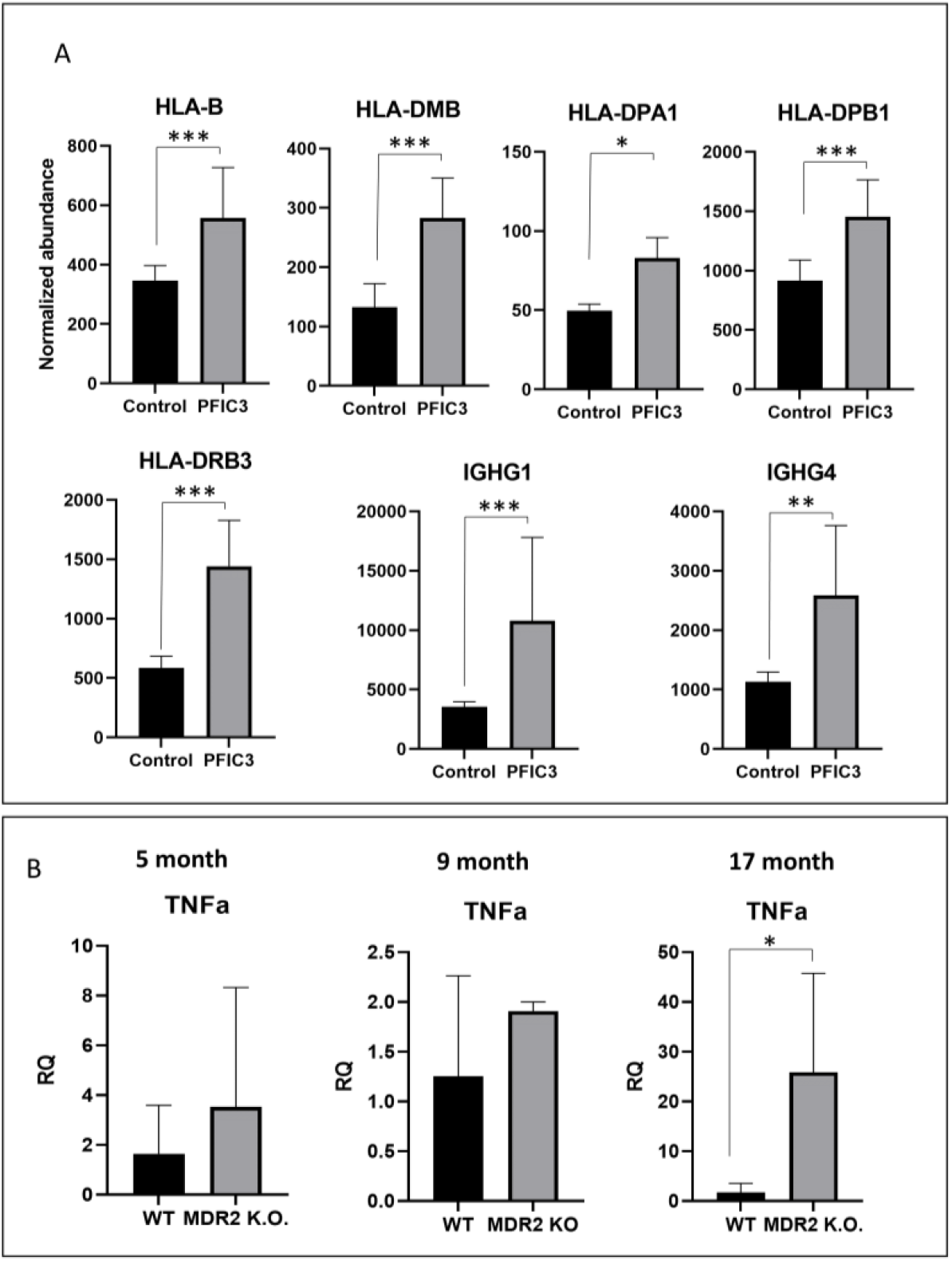
Inflammatory response of PFIC3 livers. A, up-regulated proteins associated to inflammation in PFIC3 compared to normal samples. B, overproduction of TNFa in MDR2-/-mouse compared to WT. A significant increase was detected in livers of 12-month-old mice while only trends suggested the accumulation of TNFa at 5 and 9 months. Data are the mean ± SD and statistical significance was assessed with the t-test; p values were adjusted with the Bonferroni Hochberg correction (* p<0.05; ** p<0.01; ***p<0.001).

Inflammatory reaction is a priming factor for the regenerative process required to repair a chronically injured liver^25^. Regeneration involves a phenotypic rewiring that involves a transient cell dedifferentiation and proliferation. In the cholestatic liver of PFIC3 patients we observed a marked decrease of proteins typically expressed by differentiated hepatocytes, including ADH1A and C, CYP1A2 and metallothionenines (MT) 1F, 1E, 2A and 1H (The Human Protein Atlas, www.proteinatlas.org), which are part of the detoxification machinery of the hepatocytes (Figure 4A). To investigate the mechanism underlying the down-regulation of these hepatocyte proteins, we next examined AKT and downstream GSK3 phosphorylation as their activation has been previously involved in MT down-regulation^26^. Increased phosphorylation of GSK3 and AKT was observed in the liver of 9-month-old MDR2-/-mice compared to WT littermates, which supports their mediation in the negative regulation of MTs in PFIC3 (Figure 4B).

**Figure 4.**
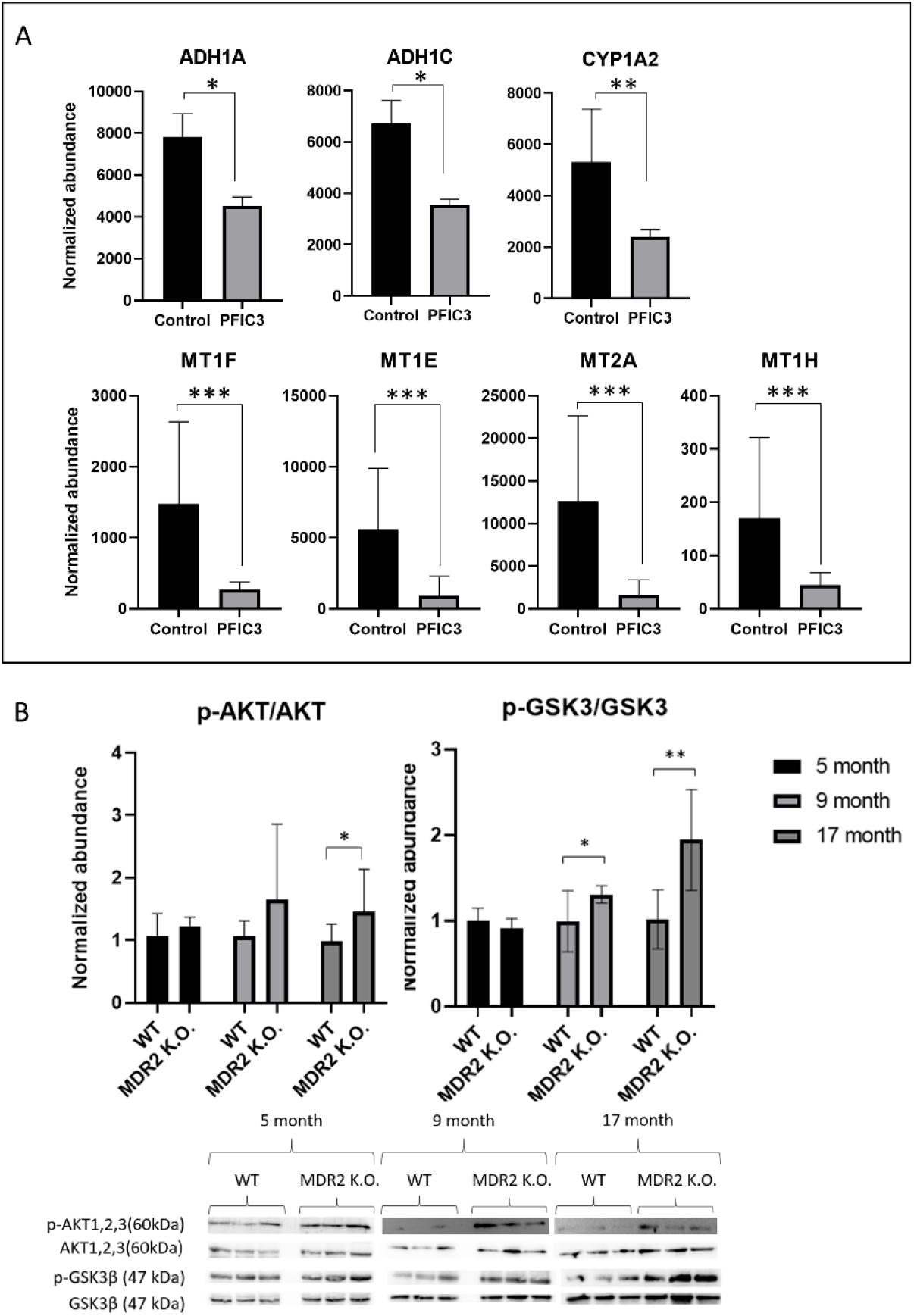
Down-regulation of liver specific proteins in PFIC patients. A, PFIC3 decrease of proteins that are mainly expressed in the liver including alcohol dehydrogenase isoforms and metallothioneins. B, Activation of AKT and GSK3 in MDR2-/-livers as a potential explanation for the observed metallothionein down-regulation in PFIC3 samples; the bar plot results from the quantification of the Western blot shown in the figure; GAPDH was used as housekeeping protein. Data are the mean ± SD and statistical significance was assessed with the t-test; p values were adjusted with the Bonferroni Hochberg correction (* p<0.05; ** p<0.01; ***p<0.001).

Activation of AKT and GSK3 also suggest the activation of cell proliferation, which is further supported by the overexpression of CDC27 and CDK5, and the activation of the HIPPO signaling pathway (Figure 5A). According to this hypothesis, activation of HIPPO pathway mediators, including CTGF, CyclinD1, FGF1 and TGFb2, was confirmed by qPCR (Figure 5B). MDR2 deficiency also induce CTGF ab Cyclin D1 but only in 17-month-old mice (Figure 5C), while 14-3-3 protein was induced only at 5 months, with no significant changes compared to WT livers at late stages (Figure 5D).

**Figure 5.**
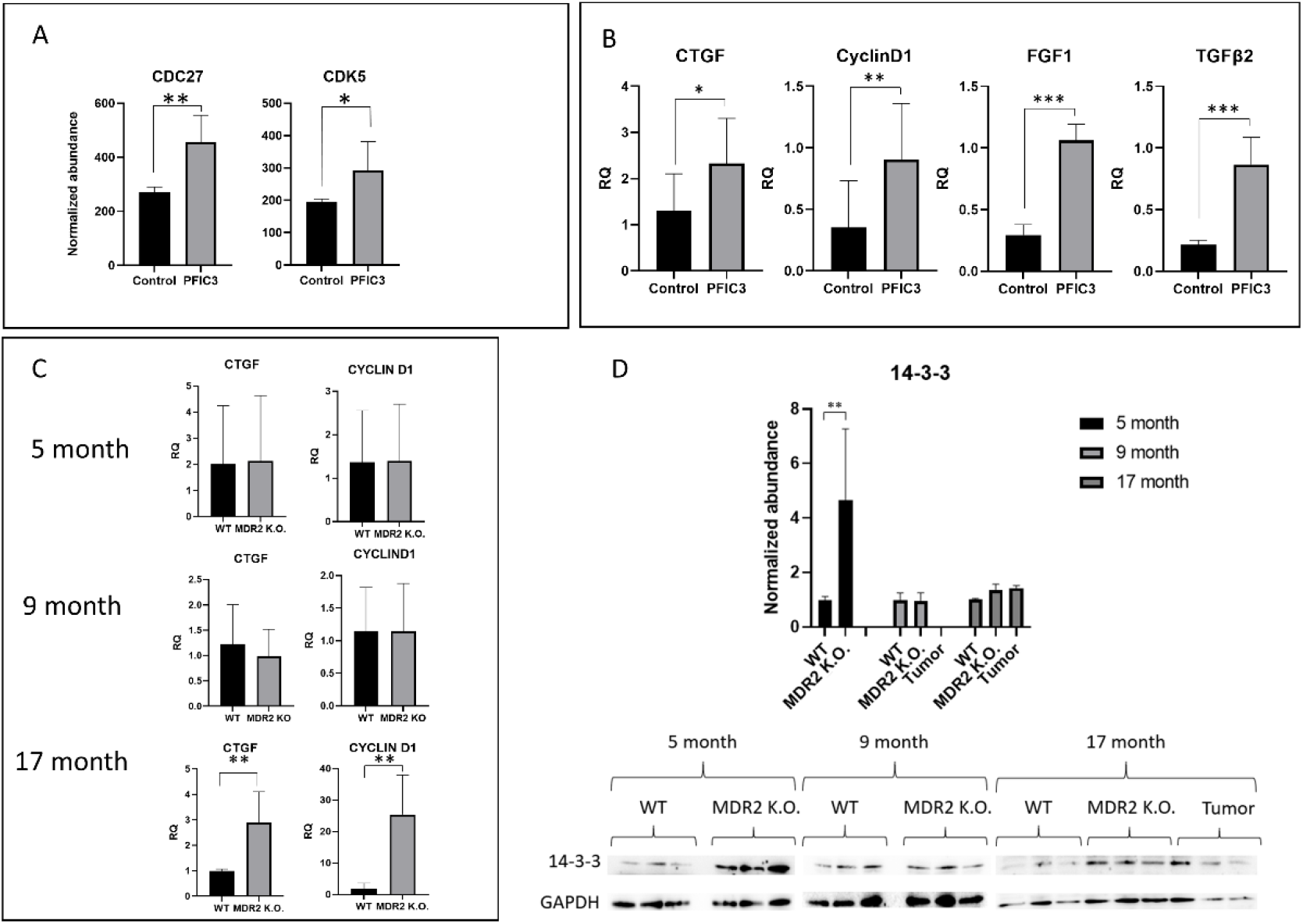
Differential proteins involved in cellular proliferation. A, Upregulation of cyclin and CDK proteins in PFIC3 compared to normal samples. B, overexpression of cell-division-associated genes in PFIC3 livers as measured by qPCR analysis. C, time course study of the expression levels of cell proliferation mediators in WT and MDR2-/-mice showing stimulation at late stages. D, up-regulation of 14-3-3 protein in MDR2-/-compared to WT livers at 5 months, suggesting the impairment of expression of HIPPO pathway target genes; Measures were done by Western blot, whose quantification is shown in the plot; GAPDH was used as housekeeping protein. Data are the mean ± SD and statistical significance was assessed with the t-test; p values were adjusted with the Bonferroni Hochberg correction (* p<0.05; ** p<0.01; ***p<0.001).

### Metabolic rewiring in PFIC3 livers

In cholestasis, the accumulation of BAs induces a compensatory response aiming to repress the expression of CYP7A1, the rate limiting step of BA synthesis from cholesterol, and CYP8B1, which is mediated by FXR-FGF15/19 activation^27,28^. As expected, we observed a decrease of CYP8B1 in PFIC3 livers (Figure 6A) that was confirmed in the liver of MDR2-/-mice, where also down-regulation of CYP7A1 was demonstrated (Figure 6B). The effect in mice was significant by 5 month and also by 8 months for CYP8B1; at later times, aging effects affecting the hepatic CYP450 levels likely overlap with the mechanisms triggered to regulate BA synthesis in cholestasis^29^. Further supporting the activation of the FGF15/19 signaling, we observed a marked decrease of PEPCK, G6PASE, and IDH3A in MDR2-/-livers of 5- and 8-month-old mice. These observations point out to an active glycolysis and to a reduced gluconeogenesis as well as an impairment of TCA, as suggested by IDH3 impairment in the MDR2-/-mouse at 5 and 9 months. At 17 months, up-regulation of these three enzymes might parallel the development of hepatocellular carcinoma (Figure 6C). Glycolysis activation (HKDC1, PFKP increase) and TCA down-regulation (ACSS2 decrease) was also observed in PFIC3 livers (Figure 7A). In parallel, the Phosphorylation of PDHA1 in serine 232 by PDHK1 inhibits the conversion of pyruvate to Acetyl-CoA^30^, blocking mitochondrial pyruvate metabolism^31^

**Figure 6.**
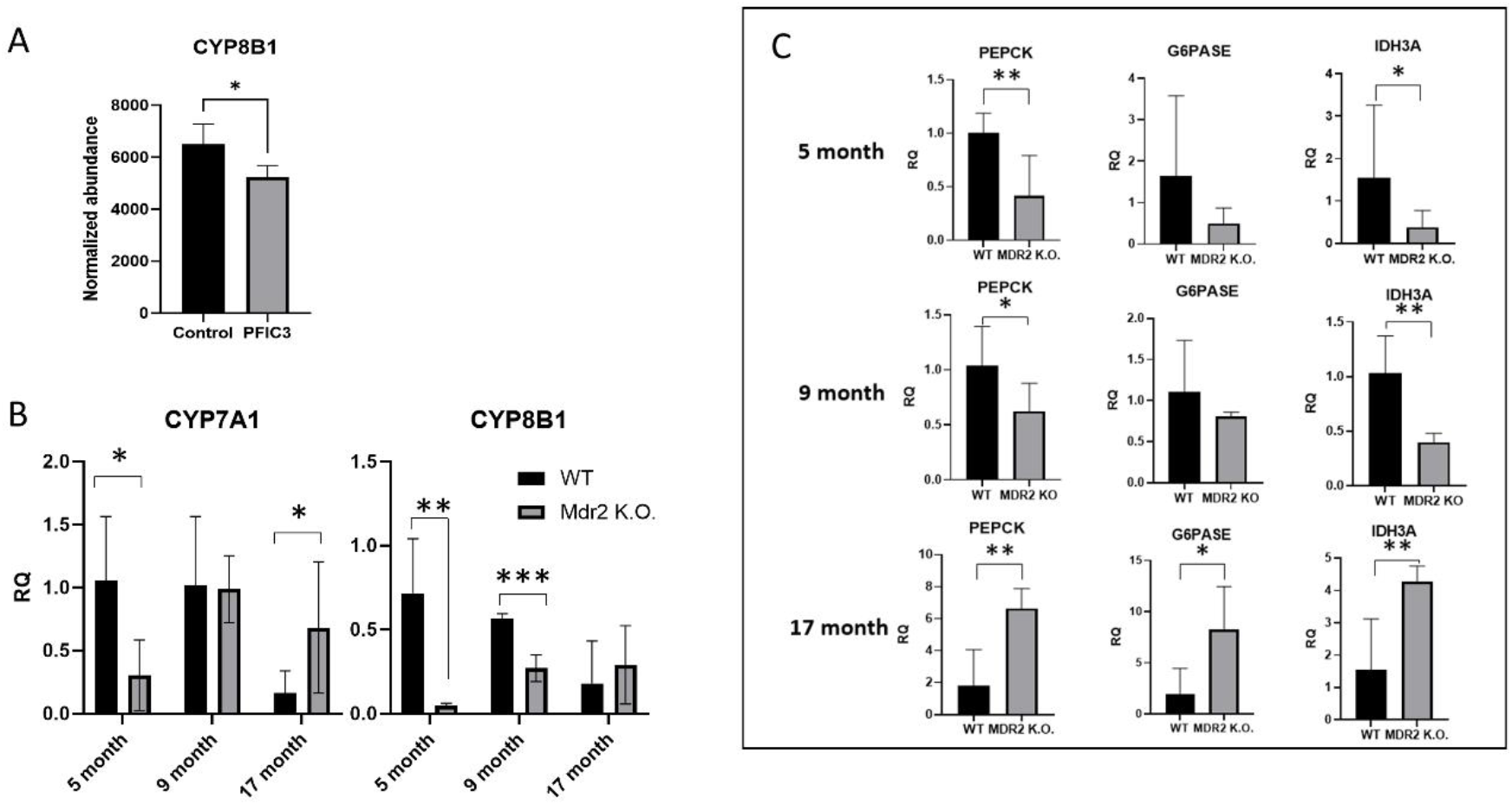
Analysis of FGF15/19 signaling. A, down-regulation of CYP8B1 in PFIC3 livers suggesting the modulation of BAs synthesis by FGF15/19. B, early down-regulation of CYP7A1 and CYP8B1 in MDR2-/-mice compared to WT liver samples as measured by qPCR. C, regulation of PEPCK, G6PASE and IDH3A in MDR2-/-mice, gene targets of FGF15/19, which suggest a metabolic rewiring in the liver associated to MDR2 deficiency. Data are the mean ± SD and statistical significance was assessed with the t-test; p values were adjusted with the Bonferroni Hochberg correction (* p<0.05; ** p<0.01).

**Figure 7.**
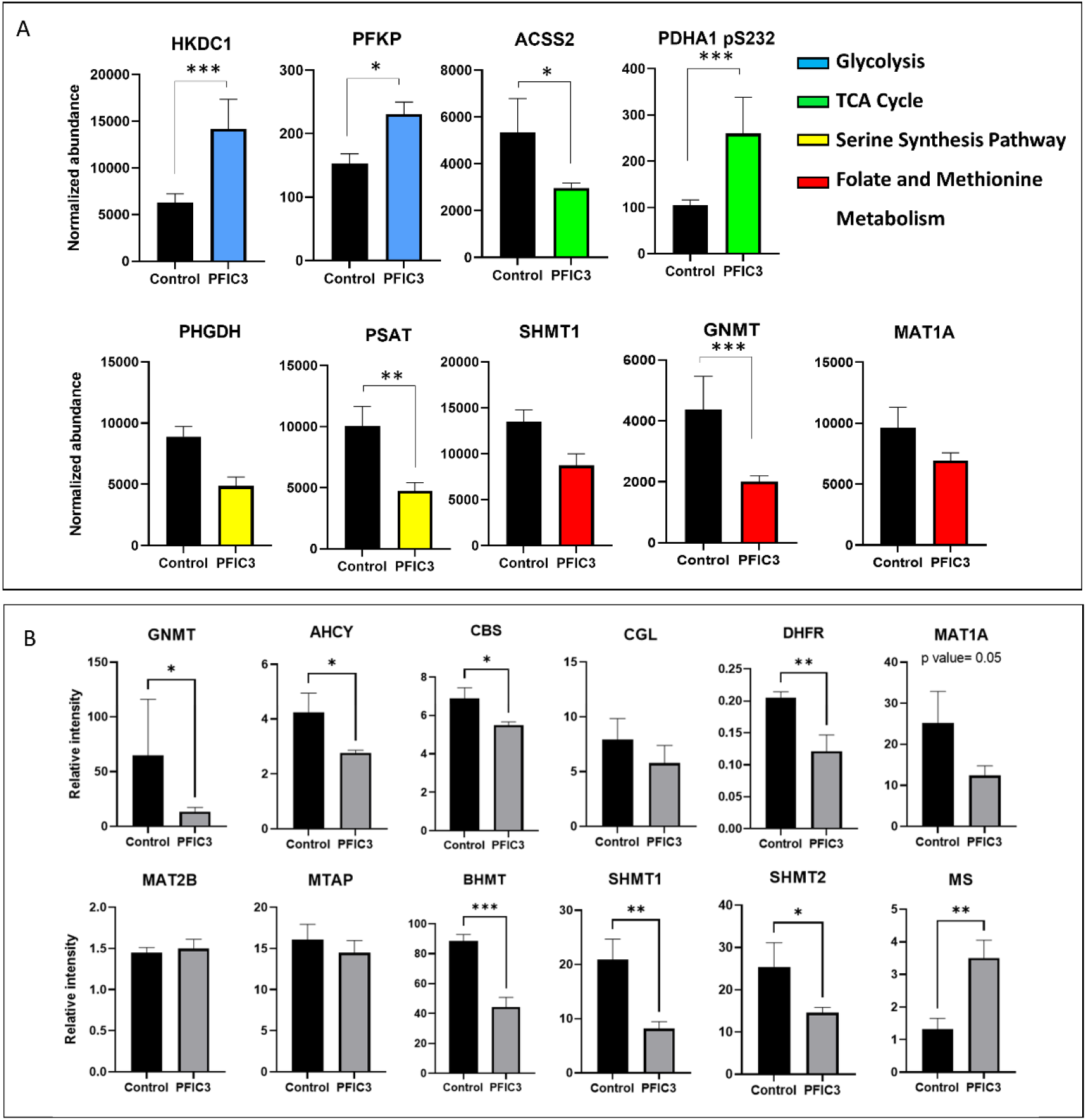
Metabolic rewiring induced in the liver by MDR3 deficiency. A, regulation of enzymes involved in the glycolysis, gluconeogenesis, TCA, OCM and the link between them in PFIC3 compared to normal liver samples. B, regulation of OCM enzymes in PFIC3 livers; measurements were done by targeted PRM. Data are the mean ± SD and statistical significance was assessed with the t-test; p values were adjusted with the Bonferroni Hochberg correction (* p<0.05; ** p<0.01; ***p<0.001).

Glycolysis is connected to one carbon metabolism (OCM) through the synthesis of serine from 3-phosphoglycerate in three consecutive steps. Two of the enzymes catalyzing this process, PSAT and PHGDH are downregulated in PFIC3 livers, suggesting a deficient OCM turnover, a hypothesis that is supported by the reduction of GNMT, and the same trend of SHMT1 and MAT1A, as evidenced by our shotgun proteomics observations (Figure 7B).

Since OCM is the link between intermediate metabolism and epigenetic regulation^32^ and is central for a normal liver function^33^, we next wanted to assess its potential impairment in the liver of PFIC3 patients. To this end, the abundance of enzymes catalyzing 12 OCM reactions was monitored using a targeted PRM approach. Most of these enzymes were significantly reduced in PFIC3 livers (Figure 7C), indicating an OCM reprograming that is in good agreement with the tissue repairing process triggered by the chronic liver injury associated to cholestasis. Interestingly, MAT2A was not detected and MAT2B showed no change with respect to normal livers, suggesting that hepatocytes in these PFIC3 patients did not undergo oncogenic transformation at this stage.

### Citoskeleton, cell adhesion and cell polarity

Cytoskeleton organization is essential in the liver organ homeostasis and disease control as it regulates cell morphology, cell polarity, intracellular trafficking, cell-cell interaction and communication with extracellular matrix components through interaction with integrins and other protein components^34^. Impaired microfilament organization has been largely recognized as one of the factors contributing to chronic liver damage as in cholestasis. We observed a downregulation of Nectin3, an adhesion protein participating at adherens junctions, as well as a decreased phosphorylation of afadin, HEPACAM, two central components of adherens junctions and cell-matrix interaction respectively (Figure 8A). Up-regulation of MYL9 expression levels, and MYH11 phosphorylation indicate a cytoskeleton rewiring (Figure 8B). To further investigate the remodeling of cytoskeleton in PFIC3, we studied the phosphorylation status MLC in MDR2-/-mice. Phosphorylation of MLC was significantly increased in MDR2 deficient livers compared to WT by 9 months of age (Figure 8C).

**Figure 8.**
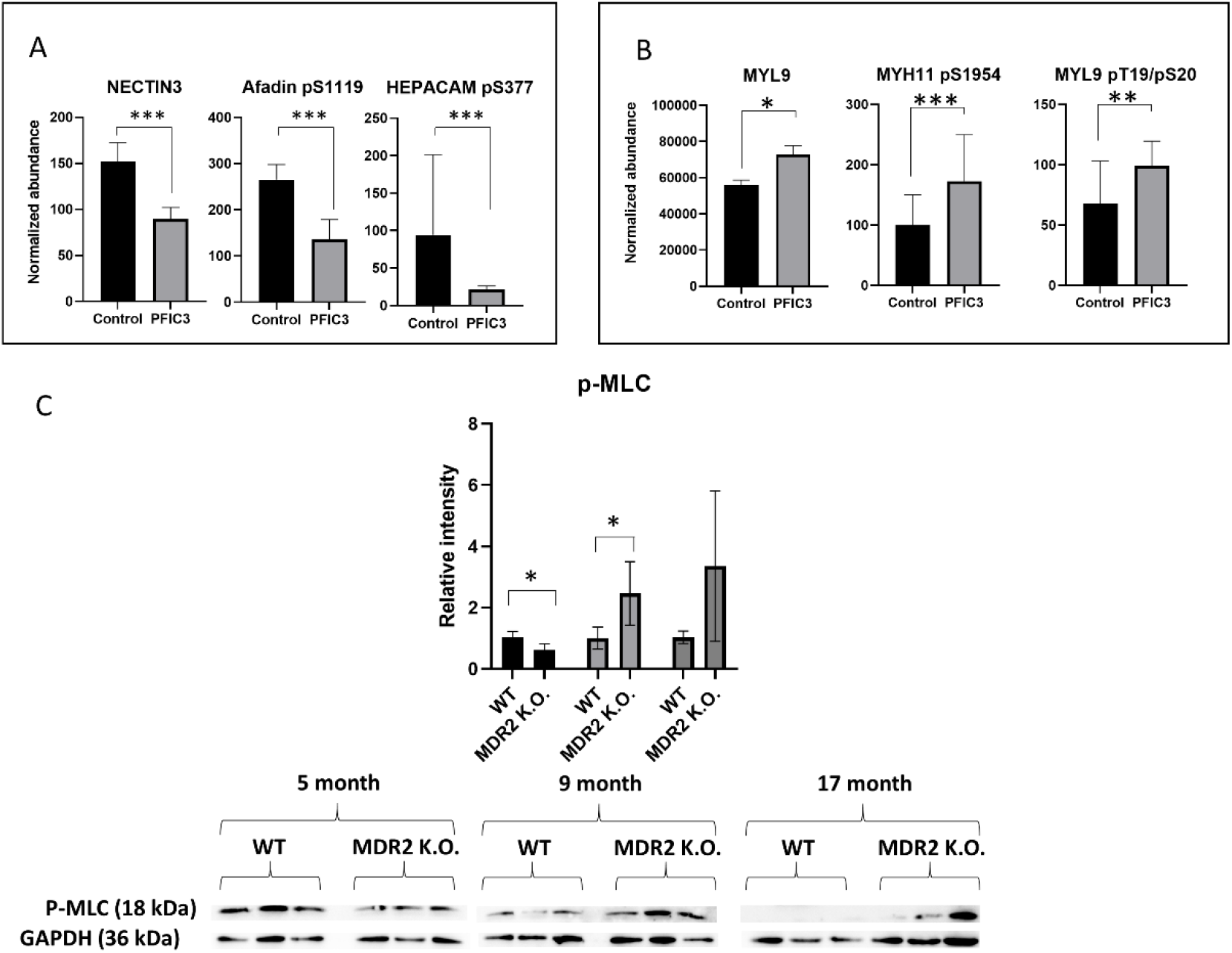
Cell adhesion and cytoskeleton alterations in PFIC3 liver samples. A, cell adhesion proteins regulated in liver samples from PFIC3 patients. B, cytoskeleton proteins regulated in PFIC3 compared to normal livers. C, regulation of MLC phosphorylation in MDR2-/-livers as measured by Western blot, whose quantification is shown in the plot; GAPDH was used as housekeeping protein. Data are the mean ± SD and statistical significance was assessed with the t-test; p values were adjusted with the Bonferroni Hochberg correction (* p<0.05; ** p<0.01; ***p<0.001).

## Discussion

PFIC represents a group of autosomal recessive disorders that are presented during the childhood as an intrahepatic cholestasis resulting from defects of bile acid synthesis and transport^3^. PFIC3 in particular is caused by mutations on ABCB4 gene encoding the phosphatidyl choline floppase MDR3 and its associated phenotype might range from a benign recurrent cholestasis, to a severe syndrome that might lead to liver failure and cancer within the first ten years of life. To date the availability of a 100% efficient therapeutic option is yet an unmet need. In this regard, the systematic analysis of the molecular basis of PFIC3 progression might provide useful information to develop new strategies for a better management of PFIC3 patients. The proteome- and phosphoproteome-wide analysis reported here provide the identification of differential protein species in PFIC3 livers that might, at least partially, explain the cellular processes triggered in the hepatocyte by the deficiency of MDR3 transporter. Based on this, an integrated rewiring of cell biology in response to the chronic cholestatic liver injury is proposed.

PFIC3 origins in *Abcb4* gene mutations that might compromise MDR3 levels, stability and activity^12^. The patients analyzed in this study are carriers of 4 different mutations on *Abcb4* gene that induce the inactivation of the resulting MDR3 variant while the levels of the protein remain unchanged with respect to individuals with normal *abcb4* alleles. However, phosphorylation at residues S666 and T667 was significantly reduced in all cases, and, to the best of our knowledge, associated to PFIC3 for the first time. These residues are located at a flexible loop between the two nucleotide binding domains but with an undetermined spatial configuration^9^. Additional studies are required to determine the kinase/phosphatase activity responsible for controlling the phosphorylation of S666 and T667 as well as the impact of these modifications on MDR3 activity. Nevertheless, as phosphorylation of MDR3 in the N-terminal region has been reported as a regulatory mechanism of MDR3 activity^19^, it is tempting to suggest that the extra negative charge represented by phosphorylation of S666 and T667 might also configure a regulatory mechanism, perhaps at the level of binding or hydrolysis of ATP.

MDR3 deficiency leads to an impaired PC transport to bile, which compromises the neutralization of the detergent effect of hydrophobic BAs, inducing a biliary epithelium injury and ultimately leads to cholestasis. In order to repair the damaged tissue, a regenerative response is orchestrated that primarily involve inflammation mediated by a CCL2-dependent proinflammatory monocyte recruitment^35^. In agreement to this, we observed up-regulation of class I HLA and immunoglobulins in PFIC3 livers. Moreover, Guillot et al also demonstrated a linked proliferative stimulation upon monocyte recruitment, which is dependent on the production of TGFb and ITGB6 and provides explanation to the observed stimulation of the HIPPO pathway in the liver of PFIC3 patients^36^, which is also confirmed in MDR2-/- mice. Moreover, it is worth noting that HIPPO signaling pathway plays a pivotal role in controlling organ size and maintaining tissue homeostasis in multiple organisms ^37^. These results provide driver proteins of the inflammatory process induced by the cellular damage associated to the presence of free hydrophobic BAs in bile and of the regenerative process initiated to repair the injured tissue, which is also related to the profibrogenic process inherent to MDR3 deficient livers^38^. Another alteration typically occurring in cholestasis is the loss of cell polarity. Our data point to impaired tight and adherens junctions. In association to this, reconfiguration of actin-myosin microfilament network in PFIC3 hepatocytes might account for the changes in morphology, dedifferentiation and compromised endosomal^39^ and Golgi-mediated^40^ intracellular trafficking. The effects associated to activation of mesenquimal, non-parenquimal, liver cells should also be taking into consideration to explain the alterations on the cytoskeleton proteome.

Concomitant to this, a significant metabolic rewiring was observed in PFIC3 livers, which might occur to fulfil the requirements of energy and biomolecules to support cell proliferation. Here we show downregulation of the expression of CYP7A1 and CYP8B1 in PFIC3 patients, which is a primary response to the accumulation of BAs in cholestasis, according to a mechanism dependent on the activation of the FXR-FGF15/19 axis^41,42^. These observations were validated in MDR2-/-mice at 5 months while the time-dependent down-regulation in the WT might be associated to ageing, as has been observed in rats^29^. Our data also support increased glycolysis, reduced gluconeogenesis, and reduction of acetyl-CoA production, which might suggest a Warburg-like effect where pyruvate is transformed into lactate as main source of energy instead of TCA cycle^43^. These observations might favor the notion of a regulated cell growth as part of a tissue regeneration process instead of an uncontrolled proliferation associated to cancer. Glycolysis is connected to one carbon metabolism through the conversion of 3-phosphoglycerate into serine in three consecutive reactions, a process that might be compromised by the consumption of 3PG in glycolysis. The glycolytic metabolic flow is further supported by the decrease of PSAT, an enzyme that catalyzes the first reaction transforming 3PG into serine. OCM serves as an integrative pathway, relating many nutrients to one another and represents the link to epigenetic regulation. Therefore, the ability to fine-tune one-carbon metabolism is essential for the maintenance of cellular homeostasis and to enable cell fate decisions^44^. In fact, changes of the expression pattern of different OCM enzymes have been described in association with chronic liver injury and HCC^33^. In the present study we described a general down-regulation of OCM enzymes in PFIC3 livers, which might facilitate cell proliferation for tissue injury repair while the maintenance of MAT2A and B levels support the regulated nature of the process, in contrast to the uncontrolled cell growth that is typically found in cancer development. It is tempting to speculate that the systematic monitoring of OCM would be a useful tool to assess the physiological status of the liver and to allow for discrimination among a controlled regenerative process and tumor progression ^45^. It has been also reported that some OCM metabolites might provide alternative therapeutic options. In this direction, administration of MTA to MDR2-/-mice reduces the production of proinflammatory and pro-fibrotic mediators and inhibits the proliferation and activation of fibrogenic cells, proving its protective role from liver injury^24^.

Overall, our results provide an integrated picture of the cellular events contributing to the onset and progression of cholestasis in PFIC3 patients. Moreover, we report the identification of driver proteins that significantly enhance our understanding of the molecular pathogenesis of this disease and suggest new options for the follow up and treatment of the afflicted patients.

## Supporting information

MRM targeted proteomics OCM method table

Heatmaps proteome and phosphoproteome

## Acknowledgments

The CNB was supported by *Comunidad de Madrid* Grants B2017/BMD-3817 and 2022/BMD-7232. Severo Ochoa Project SEV 2017-0712. Intramural CSIC PIE/COVID-19 projects 202020E079 and 202020E108. MICIN PID2021-127496NB-100. This research work was also funded by the European Commission—NextGenerationEU (Regulation EU 2020/2094), through CSIC’s Global Health Platform (PTI Salud Global) and Conexión Cancer.

