## Supplementary material for "Molecular basis of Progressive Familial Intrahepatic Cholestasis 3. A proteomics study": MRM targeted proteomics OCM method table

| **Protein** | **Gene** | Code | Peptide | **Transitions** |
|---|---|---|---|---|
| **Glycine-N-Methtyltransferase** | GNMT | Q14749 | VWQLYIGDTR++<br>AWLLGLLR ++<br>AGGLLVIDHR ++ | y8+, y7+, 76+, y2+<br>y6+, y5+, y4+ b2+<br>y6+, y5+, y4+, y2+ |
| **S-adenosyl homocysteinase** | AHCY | P23526 | VADIGLAAWGR++<br>VPAINVNDSVTK ++<br>VAVVAGYGDVGK ++ | y9+, y7+, y5+, b3+<br>y9+, y8+, y6+<br>y9+, y8+, y7+, b3+ |
| **Cystathionine beta synthase** | CBS | P35520 | ILPDILK++<br>ALGAEIVR ++<br>SNDEEAFTFAR ++ | y5+, y4+, y3+, b2+<br>y6+, y5+, y4+<br>y7+, y6+, y5+ |
| **Cystathionine gamma-lyase** | CGL | P32929 | ISFVDCSK ++<br>LLEAAITPETK ++ | y7+, y6+, y5+, y4+<br>y9+, y5+, y4+ |
| **Dihydrofolate Reductase** | DHFR | P00374 | NGDLPWPPLR++<br>LTEQPELANK ++ | y6+, y5+, y4+, b3+<br>y8+, y7+, y6+ |
| **S-adenosylmethionine synthase I** | MAT1A | Q00266 | SGLLPWLRPDSK ++<br>FVIGGPQGDAGVTGR ++ | y7+, y6+, y2+, b3+<br>y12+, y10+, y8+ |
| **Methionine Adenosyltransferase II beta** | MAT2B | Q9NZL9 | VLVTGATGLLGR ++<br>AVLENNLGAAVLR ++ | y10+, y9+, y8+, b2+<br>y10+, y9+, y7+, y6+ |
| **Methylthioadenosine phosphorylase** | MTAP | Q13126 | IGIIGGTGLDDPEILEGR++ | y8+, y7+, y5+, y4+, |
| **Betaine homocysteine methyltransferase** | BHMT | Q93088 | ISGQEVNEAACDIAR ++<br>EAYNLGVR ++<br>AIAEELAPER ++ | y9+, y7+, y6+<br>y5+, y4+, y3+<br>y8+, y4+, y3+ |
| **Serine hydroxymethyl transferase I** | GLYC | P34896 | AVLEALGSCLNNK ++<br>LGTPALTSR ++ | y9+, y8+, y7+<br>y6+, y5, y4+ |
| **Serine hydroxymethyl transferase II** | GLYM | P34897 | TGLIDYNQLALTAR++<br>EYSLQVLK++<br>SAITPGGLR ++ | y10+, y9+, y8+, b3+<br>y6+, y5, y4+<br>y7+, y6+, y5+ |
| **Methionine synthase** | METH | Q99707 | YSAPVIHVLDASK ++ | y8+, y7+, y6+ |
