## Supplementary materials and methods for "Molecular basis of Progressive Familial Intrahepatic Cholestasis 3. A proteomics study"

*Biological samples*

Human tissue samples. Liver tissue fragments from the explanted livers of PFIC3 patients (N=4) and from the liver graft of living donors (N=4) were obtained at the time of liver transplantation and stored frozen in RNAlater solution (Invitrogen). ABCB4 pathogenic mutations were documented in all PFIC3 by genetic analysis. Liver transplant were performed between 6 and 9 years old. All surgical procedures were conducted in Hospital Universitario La Paz. Liver samples were kept at the institutional biobank. Written informed consent was obtained from patient's legal guardians and graft donors. Study protocols conformed to the principles stated in the Declaration of Helsinki and as such they were approved by the Clinical Research Ethics Committee of Hospital Universitario La Paz.

*Mdr2* -/- and wild type mice liver. Age: 5 months (WT: n=3; *Mdr2*: -/- n=3), 9 months (WT: n=3; *Mdr2* -/-: n=3), and 17 months (WT: n=3; *Mdr2* -/- non-tumor: n=3; *Mdr2* -/- tumor: n=3),

*Lysis and protein extraction*

Each liver sample (human and mice) was thawed and disrupted mechanically using a Potter–Elvehjem homogenizer in lysis buffer containing 5% SDS (Sodium dodecyl sulfate) (Sigma-Aldrich), 100 mM triethylamonium bicarbonate (Thermo Fisher Scientific) and a protease/phosphatase inhibitor cocktail (Thermo Fisher Scientific). For thorough homogenization and removing DNA, samples were sonicated by micro tip probe ultrasonication for 1 min on UP50H ultrasonic lab homogenizer (Hielscher Ultrasonics). After centrifugation at 10,000 × *g* for 5 min, the protein concentration of the saved supernatant was measured using the Pierce 660-nm Protein Assay adding IDCR (Ion detergent compatibility reagent) (ThermoFisher Scientific). Supernatant containing the proteins were stored at -80ºC until further analysis by LC-MS and western blot.

*Liquid chromatography and mass spectrometry analysis (LC-ESI-MS/MS)*

Liver proteins were reduced and alkylated by adding 5 mM TCEP (tris(2-carboxyethyl) phosphine) and 10 mM chloroacetamide for 30 minutes at 60ºC before digestion into peptides. Protein digestion was performed in the S-Trap filter (Protifi, Huntington, NY, USA). 80 µg of protein of each sample was diluted to 40 µL with 5% SDS, 1.2% phosphoric acid and seven volumes of binding buffer (90% methanol; 100 mM TEAB) and then loaded to a S-Trap filter. Then the filter was washed 3 times with 150 µL of binding buffer. Finally, 4 µg of Pierce MS-grade trypsin (Thermo-Fisher Scientific) in 20 μL of a 100 mM TEAB solution was added to each sample in a ratio 1:20 (trypsin/protein) and to digestion was performed in a wet chamber at 37°C overnight. To elute peptides, two step-wise buffers were applied: first, 40 μL of 25 mM TEAB and then, 40 μL of 80% acetonitrile and 0.2% formic acid in $H_2O$, separated by a 2 min centrifugation at 3000 x g in each case. Peptide amount was determined using Qubit 2.0 Fluorometer (Thermo Fisher Scientific). Eluted peptides were pooled and vacuum centrifuged to dryness in a Speed Vac concentrator (Eppendorf).

The resulting peptides were subsequently labelled using TMT-11plex Isobaric Mass Tagging Kit (Thermo Scientific, Rockford, IL, USA) according to the manufacturer's instructions as follows: 126: WT-1; 127N: PFIC3-1; 127C: WT-2; 128N: PFIC3-2; 128C: PFIC3-3; 129N: WT-3; 129C: PFIC3-4; 130N: WT-4; 130C: PFIC3-3 replicate; 131N: WT-1 replicate; 131C: internal standard pool). Labelled peptides were used for the analysis of the global proteome and the phosphoproteome.

For deep analysis of the global proteome, high pH reversed-phase peptide prefractionation was performed with sulfonated divinylbenzene (CDS Empore™ SDB-RPS, Sigma-Aldrich) using a step gradient of increasing acetonitrile concentrations (0-60% ACN) in a high pH elution solution (10 mM ammonium formate ($NH_4HCO_2$), pH 10.0). Eluted peptides were collected into 10 different fractions by centrifugation at 850 xg during 3 minutes. The peptide fractions were dried and stored at −20 °C until the LC−MS analysis.

In parallel, phosphopeptide enrichment was performed using $TiO_2$ columns and cleaning of the sample was performed with Oligo R3 reverse-phase before LC-MS analysis.

Peptides were solubilized in 2% acetonitrile (ACN) and 0.1% formic acid (FA), then peptide concentration was determined by Qubit 2.0 Fluorometric Quantitation (Thermo Fisher Scientific). 1 µg aliquot of each fraction in an injection volume of 5 µl was analyzed by nano LC ESI-MS/MS (Liquid Chromatography Electrospray Ionization Tandem Mass Spectrometry) in an Ultimate 3000 nano HPLC system (Thermo Fisher Scientific) coupled to an Orbitrap Exploris 240 (Thermo Fisher Scientific). Peptides were loaded onto a 50 cm × 75 μm Easy-spray PepMap C18 analytical column at 45°C and were separated at a flow rate of 300 nL/min using a 90 min gradient ranging from 2 % to 95 % mobile phase B (mobile phase A: 0.1% FA; mobile phase B: 80 % ACN in 0.1% FA).

For fractions corresponding to the total proteome, data acquisition was performed using a data-dependent top-20 method, in full scan positive mode, scanning 375 to 1200 m/z.

Survey scans were acquired at a resolution of 120,000 at m/z 200, with Normalized Automatic Gain Control (AGC) target (%) of 300 and a maximum injection time (IT) in AUTO. The top 20 most intense ions from each MS1 scan were selected and fragmented via Higher-energy collisional dissociation (HCD). Resolution for HCD spectra was set to 45,000 at m/z 100, with AGC target of 50 and a maximum ion injection time in AUTO. Isolation of precursors was performed with a window of 2 m/z, exclusion duration (s) of 45 and the HCD collision energy was 30. Precursor ions with single, unassigned, or six and higher charge states from fragmentation selection were excluded.

For the phosphopeptides enriched fraction, data acquisition was performed using a data-dependent top-20 method, in full scan positive mode, scanning 375 to 1200 m/z. Survey scans were acquired at a resolution of 60,000 at m/z 100, with Normalized Automatic Gain Control (AGC) target (%) of 300 and a maximum injection time (IT) in AUTO. The top 20 most intense ions from each MS1 scan were selected and fragmented via Higher-energy collisional dissociation (HCD). Resolution for HCD spectra was set to 45,000 at m/z 100, with AGC target of 200 and a maximum ion injection time in AUTO. Isolation of precursors was performed with a window of 2 m/z, exclusion duration (s) of 10 and the HCD collision energy was 30. Precursor ions with single, unassigned, or six and higher charge states from fragmentation selection were excluded.

*Proteomics data analysis*

Raw instrument files were processed using Proteome Discoverer (PD) version 2.4 (Thermo Fisher Scientific). MS2 spectra were searched using four search engines (Mascot *(*v2.7.0), MsAmanda (v2.0), MsFragger (v3.1.1) and Sequest HT and a target/decoy database built from sequences in the human proteome at Uniprot Knowledgebase (9606rev_20210219). All searches were configured with dynamic modifications pyrrolidone from Q (-17.027 Da) and oxidation of methionine residues (+15.9949 Da) and static modification for TMT reagents (+229.163 Da) on lysine and N-term of the peptide, and carbamidomethyl (+57.021 Da) on cysteine. For the phosphopeptides search, a dynamic modification of addition of phosphate group was included for serine, threonine and tyrosine residues (+79.966). Trypsin cleavage was configured (max 2 missed cleavages). The peptide precursor mass tolerance was 10 ppm, and MS/MS tolerance was 0.02 Da. The false discovery rate (FDR) for proteins, peptides, and peptide spectral matches (PSMs) peptides were kept at 1%. The quantification values for proteins were calculated using the abundance of total peptide for the identification of differentially expressed proteins. In this case, the peptide group abundances were summed for each sample and the maximum sum for all files was determined.

To calculate the p-values and adjusted p-values for quantification results, the "*t-test background based*" statistical method was used. Those proteins differentially expressed with an adjusted p-value >= 0.05 were considered as significant. A list of differentially expressed and phosphorylated proteins was used to perform an Ingenuity Pathway Analysis (IPA).

*Monitoring One-carbon metabolism by targeted proteomics (MRM)*

For monitoring One-carbon metabolism in liver samples, 12 participating enzymes were monitored by multiple reaction monitoring (MRM) targeted proteomics: Adenosylhomocysteinase (AHCY), Betaine-homocysteine S-methyltransferase (BHMT), Cystathionine β-synthase (CBS), Cystathionine γ-lyase (CGL), Dihydrofolate reductase (DHFR), Glycine N-methyltransferase (GNMT), Methionine adenosyltransferase 1A/2B (MAT1A/MAT2B), Methionine synthase (MS), S-methyl-thioadenosine phosphorylase (MTAP), Serine hydroxymethyltransferase 1/2 (SHMT1/2). The selection of proteotypic peptides for targeted proteomics monitoring combined information from public databases as SRMAtlas (http://www.srmatlas.org) and PeptideAtlas (http://www.peptideatlas.org), as well as in-house generated human liver proteome library as previously detailed in *Journal of physiology and Biochemistry*, Guerrero et al, 2022. 27 peptides were selected according to the following criteria: no missed cleavages, peptides with Met, Trp, or other amino acids that might be modified either in the cellular environment or during the analysis were avoided if alternative peptides were available (Supplementary table). 3-4 most intense transitions according to the spectral library were selected for precursor monitoring. Retention time of each peptide in the analytical column was determined according to a first sample injection were all peptides included in the MRM method were monitored along the gradient (non-scheduled method) and, then, a scheduled method (5 minutes window) was developed. Heavy version containing 13C and 15N lysine (+8 Da) and arginine (+10 Da) were synthetized using standard F-moc chemistry. The quantification was calculated as the ratio (light/heavy) total area resulting from the sum of all the transitions of each peptide. MRM analyses were performed with 1 μg total peptide amount as determined on a Qubit 2.0 fluorimeter. Peptides were loaded into a C18 PepMap trapping column (5-μm particle size, 100 μm I.D. × 5 cm; ThermoFisher Scientific) at 2 μL/min flow rate of 0.1% FA and then separated on a C18 column (3-μm particle size 120 Å pore size, 75 μm

I.D. × 15.2 cm) (Nanoseparations, Nieuwkoop, The Netherlands). Elution was achieved with a 60-min stepwise gradient of ACN in 0.1% FA: from 2 to 40% ACN in 42 min, 40 to 95% ACN in 7 min, and 3 min in 95% ACN before re-equilibration in 2% ACN. Peptide separation was performed at 300 nL/min and 40 °C. MS/MS analyses were done on a 5500 QTRAP triple-quadrupole mass spectrometer setting a dwell time of 20 ms (for non-scheduled methods) and a declustering potential of 80 V. Raw MRM data files were analyzed with Skyline software (v21.2.0.568), and the peak selection in the chromatograms for each peptide was manually curated. Transitions showing some interference in peak area were excluded. The intensity area of each peak was automatically calculated by the software considering the value as the ratio unlabeled/SIL precursors. Statistical analysis and graphical representation of data were performed using GraphPad Prism Software v9.3.1 was used. Statistical significance is represented as *: p-value <0.05, **: p-value <0.005, ***: p-value <0.001 according to t-test analysis results.

*Western blot*

20 µg of protein were resolved in loading buffer (10% glycerol, 2 % SDS, 5% β mercaptoethanol, 0.062 M Tris-HCl, 0.002 % bromophenol blue). Electrophoresis was performed in 12% acrylamide at constant amperage (20 mA). The, proteins were and transferred to 0.2 µm nitrocellulose membranes. Membranes were blocked with 1 × TBS/0.05% Tween containing 5% milk or 5% bovine seroalbumin (BSA) according to the instructions of the antibodies' manufacturer. Membranes were then incubated with primary antibodies at 4 °C overnight in 1 × TBS/0.05% Tween containing 5% BSA/milk, and then in horseradish peroxidase (HRP)-conjugated secondary antibodies (Dako) at room temperature for 1 hour and then developed with WesternBright ECL HRP substrate (Advansta). The following primary antibodies were used: anti-AKT1, 2, 3 (Abcam Ref ab283852) anti-phosho-AKT (Ser472 + Ser473 + Ser474) (Abcam Ref ab283852), anti-mTOR (Abcam Ref ab283852), anti-ERK (Cell signaling, Ref 4695), anti-phospho-ERK (Thr202/Tyr204) (Cell signaling, Ref 4370), anti-GSK3β (Genetex Ref GTX635816) anti-phospho-GSK3 β (Ser9) (Genetex Ref GTX00971), anti-phospho-Myosin Light Chain (Ser19) (Cell Signaling Ref 3671), anti-Myosin Heavy Chain II A (Biolegend Ref 909802), anti-14-3-3 (Genetex Ref GTX133736). Statistical analysis of the results was performed using R v.4.1.3. For comparisons between two groups, the t-test function of R Base package was used. Differences were considered significant when p value <0.05.

*RT-qPCR*

For RNA extraction, liver tissue samples were homogenized using a Pellet Pestle in buffer provided by RNeasy kit (Qiagen) according to manufacturer instructions. RNA concentration was determined using a Nanodrop ND-1000 (Thermo Fisher Scientific). 1 µg of RNA of each sample was retrotranscribed using the Applied Biosystems Kit RNA-to-cDNA. Quantitative PCR was then performed using EvaGreen Master Mix (Solis BioDyne) detected by the QuantStudio 5 and analyzed using QuantStudio Software v1.5.2 (Applied Biosystems). Primer sequences used for qRT-PCR are shown in table X. Results were normalized to the expression of GAPDH and presented as fold induction with respect to the wild type samples according to the $2^{-\Delta\Delta CT}$ method. Statistical analysis of the results was performed using R v.4.1.3. For comparisons between two groups, the t-test function of R Base package was used. Differences were considered significant when p value <0.05.

| Species | Gene name | Forward and reverse primer sequence (5' → 3') |
|---|---|---|
| *Mouse* | *Ctgf* | TGCGAAGCTGACCTGGAGGAAA |
| | | CCGCAGAACTTAGCCCTGTATG |
| | *Cyclin D1* | GCAGAAGGAGATTGTGCCATCC |
| | | AGGAAGCGGTCCAGGTAGTTCA |
| | *CYP7A1* | AGCAACTAAACAACCTGCCAGTACTA |
| | | GTCCGGATATTCAAGGATGCA |
| | *CYP8B1* | CATGAAGGCTGTGCGTGAGGAA |
| | | CATCACGCTGTCCAACACTGGA |
| | *GAPDH* | CAATGAATACGGCTACAGCAAC |
| | | AGGGAGATGCTCAGTGTTGG |
| | *G6Pase* | GTGGCAGTGGTCGGAGACT |
| | | ACGGGCGTTGTCCAAAC |
| | *Idh3a* | TCGTCACCATCCGAGAGAAC |
| | | GCACAACCCCATCAACGAT |
| | *Pepck* | CACCATCACCTCCTGGAAGA |
| | | GGGTGCAGAATCTCGAGTTG |
| | *TNFα* | TGCCTATGTCTCAGCCTCTT |
| | | GAGGCCATTTGGGAACTTCT |
| *Human* | *BCL2* | ATCGCCCTGTGGATGACTGAGT |
| | | GCCAGGAGAAATCAAACAGAGGC |
| | *Ctgf* | CTTGCGAAGCTGACCTGGAAGA |
| | | CCGTCGGTACATACTCCACAGA |
| | *Cyclin D1* | TCTACACCGACAACTCCATCCG |
| | | TCTGGCATTTTGGAGAGGAAGTG |
| | *FGF1* | ATGGCACAGTGGATGGGACAAG |
| | | TAAAAGCCCGTCGGTGTCCATG |
| | *TGFβ2* | AAGAAGCGTGCTTTGGATGCGG |
| | | ATGCTCCAGCACAGAAGTTGGC |
