## Supplementary figures and images for "Molecular basis of Progressive Familial Intrahepatic Cholestasis 3. A proteomics study"

### Heatmaps proteome and phosphoproteome

#### Proteome

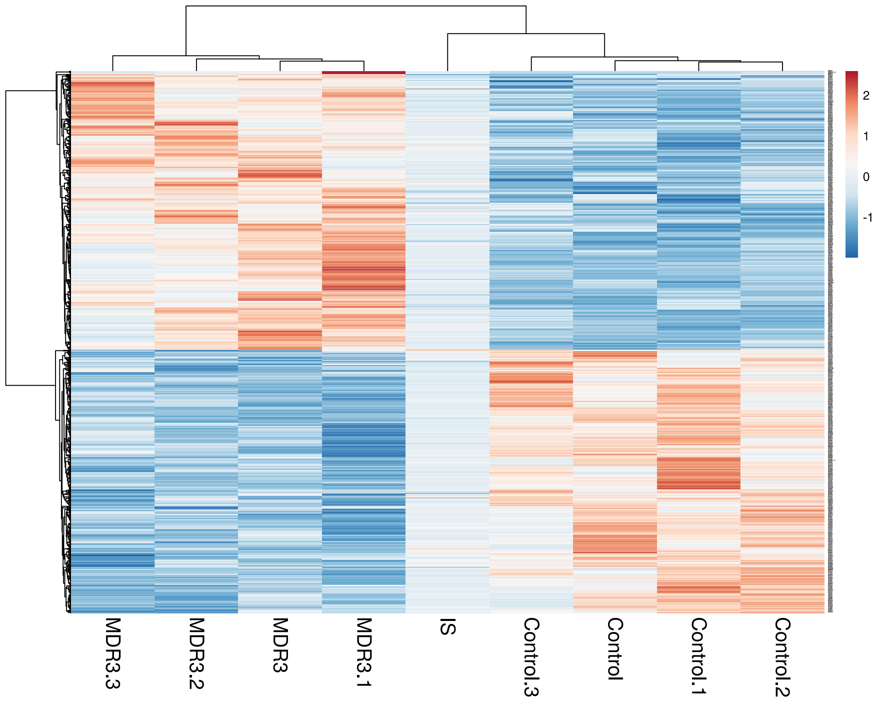


#### Phosphoproteome

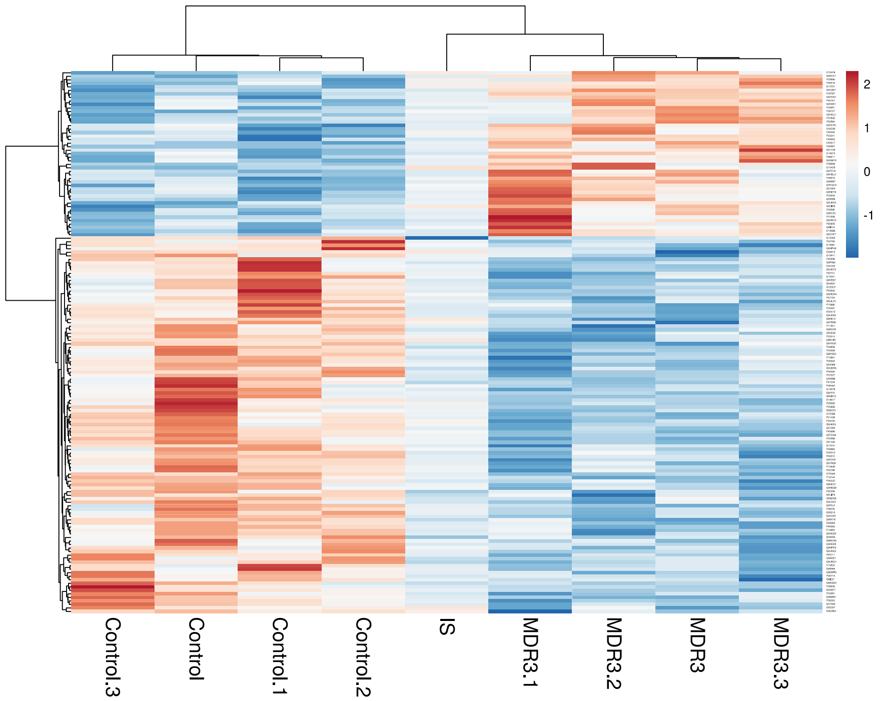


**PFIC3**

t.test adj p value < 0.05

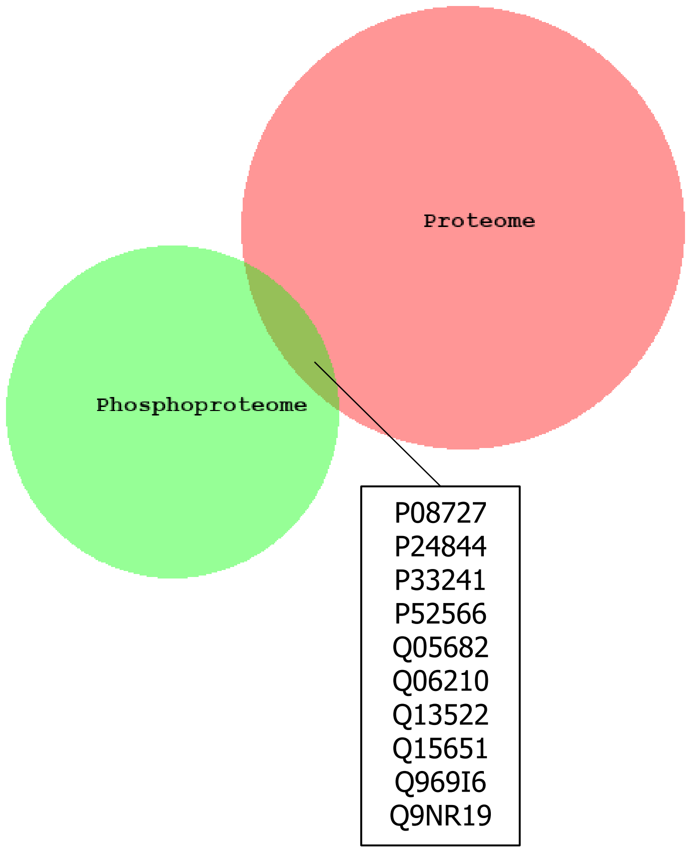
